# Strategy for Efficient Importing of NAD Analogs

**DOI:** 10.1101/480269

**Authors:** Lei Wang, Dayu Yu

## Abstract

It is difficult to study and manipulate the functions of NAD(P). NAD analogs can enhance our ability to understand signaling pathways mediated by NAD. But the application is limited. By constructing an efficient importing system will stimulate developing and screening of functional analogs.

## 1. Introduction

Recently two NAD manipulation strategies were developed to transport NAD directly across the cytoplasmic membrane, one was transporting the cofactor by NAD transporter, another was making cells permeable for the cofactors by proper treatment with certain organic solvent [2,5,6]. The NAD transporter could adjust intracellular NAD level in a wide range and the permeabilized cells realized high product concentration with high cofactor total turnover number [6,7]. Since NAD was added directly by the two strategies, the degradation characters of NAD should be considered for more efficient and precise manipulation.

To improve the efficacy of redox biocatalysts, it is essential to maintain the stability of nicotinamide cofactors, for which it is attractive to block degradation pathways for NAD(H). While the biosynthesis of NAD(H) has been well studied, it is less understood how NAD(H) are degraded. In *E. coli*, only cytoplasmic NudC has been known as NAD(H) pyrophosphatase, which hydrolyzes NAD to nicotinamide mononucleotide (NMN) and adenosine monophosphate (AMP). NudC had low catalytic efficiencies for NADH (0.03 μM^−1^ s^−1^) and NAD (2.9×10^−4^ μM^−1^ s^−1^) [8].

As NAD was impermeable, its metabolism could be studied as intracellular metabolism and extracellular metabolism respectively[8]. Study and compare intra– and extracellular metabolism of NAD might help improving NAD manipulation strategies and revealing how *E. coli* maintained its cofactor pool. Most reported NAD pyrophosphatases localized outside the cytoplasm[9]. Among the prokaryotic NAD pyrophosphatases, NadN from *Haemophilus influenzae* localized outside the cytoplasm and had two activities: it hydrolyzes NAD to NMN and AMP by NAD pyrophosphatase activity, then NMN and AMP are hydrolyzed to NR and Ado respectively by 5’–nucleotidase activity[10].

In this report we demonstrated that UshA was a major periplasmic enzyme for NAD degradation in *E. coli*. By constructing an efficient importing system will stimulate developing and screening of functional analogs.

## 2. Materials and methods

### 2.1 Regents and strains

EasyPfu DNA polymerase was from TransGen Biotech (Beijing, China). PrimeSTAR HS DNA polymerase and DpnI were from TaKaRa (Dalian, China). All the primers were synthesized by Invitrogen. All reagents and enzyme substrates were from Bonuo (Dalian, China). Yeast ADH II was from Sigma. NMN was prepared as reported previously [10].

*E. coli* BW25113 (*rrnB3,* Δ*lacZ4787, hsdR514,* Δ*(araBAD)567,* Δ*(rhaBAD)568, rph–1*) and its *ushA* deletion mutant *E. coli* JW0469 (BW25113/*ushA::kan*) was obtained from the Coli Genetic Stock Center [11]. *E. coli* WL001 was constructed by transformation of *E. coli* JW0469 with the UshA expression plasmid pA–ushA.

### 2.2 NAD degradation activity of whole cells

*E. coli* BW25113 and JW0469 cells were grown aerobically at 37 °C, 200 rpm overnight in LB liquid medium (10 g tryptone, 5 g yeast extract, 10 g NaCl per liter). For JW0469 cells, 50 μg/mL kanamycin was applied in the medium. *E. coli* WL001 cells were prepared at 30 °C, 200 rpm for 36 h in LB liquid medium supplemented with 100 μg/mL ampicillin and 1 mM isopropyl thiogalactoside (IPTG).

The cells were harvested by centrifugation at 2000 ×g for 6 min at 4 °C, washed twice and suspended with 50 mM Tris–Cl (pH 7.5) to an optical density at 600 nm (OD_600nm_) of 22, which is approximately 7.3 g cell dry weight per liter (g_CDW_/L) [2]. The cells were stored at –80 °C before use.

NAD degradation activity of whole cells was assayed by measuring extracellular NAD concentration drop discontinuously. The reactions were carried out in the presence of 50 mM Tris–Cl (pH 7.5), 1 mM NAD, 50 mM MgCl_2_, and 3.7 g_CDW_/L *E. coli* BW25113 or JW0469 cells. *E. coli* WL001 samples were assayed at a final concentration of 53 mg_CDW_/L.

The mixture was incubated at 37 °C for 30 min and quenched by adding 0.1 volume of 2 M HCl with further incubation at 50 °C for 10 min. The mixtures were neutralized by equal volume of 0.1 M NaOH, centrifuged at 12,000 ×*g* for 10 min, and the supernatants were transferred to new tubes, stored at –20 °C before NAD concentration analysis.

NAD concentration were determined by cofactor recycling assay [6]. One unit of NAD degradation activity (1 U) was defined as hydrolyzing 1 nmol NAD per hour.

### 2.3 Thin layer chromatography (TLC) analysis

Enzymatic products of UshA were assayed by TLC on silica gel according to a previous procedure [13, 14]. Reactions were done at 37 °C for 2 h in 50 μL of 50 mM Tris–Cl (pH 7.5) contained 5 mM MgCl_2_, 10 mM NAD, 0.1 mg/mL BSA and appropriate amounts of UshA. Reactions were terminated by adding 5 μL of 2 M HCl, and centrifuged at 12,000 × *g* for 10 min. Spotted 1.5 μL of samples onto TLC plates, developed in a water/acetic acid /n–butanol (5: 3: 12; vol/vol/vol) mixture, and visualized under UV at 254 nm.

### 2.4 Stability of NAD supplied to permeabilized cell

Permeabilized cells of BW25113 and JW0469 were prepared according to a previous process [5]. Briefly, frozen *E. coli* cells were thawed in a water bath at room temperature for 10 min, added EDTA to 5 mM and toluene to 1% (vol/vol), and incubated at 200 rpm, 30 °C for 30 min and then at 4 °C for 1 h. Cells were recovered by centrifugation at 2000 ×g for 6 min, washed twice and suspended with Tris–Cl (50 mM, pH 7.5). Permeabilized cells were used immediately after preparation.

To determine the half–life time of exogenous NAD in the presence of permeabilized cells, a reaction mixture with 1 mM NAD, 50 mM MgCl_2_, and 3.7 g_CDW_/L permeabilized cells were incubated at 37 °C, pH 7.5. Aliquots of 90 μL of reaction mixture were taken at 0 h, 0.5 h, 1 h, 1.5 h, 2 h, 3 h and 4 h, and NAD concentration was determined.

Half–life time of exogenous NAD was calculated according to the equation: logN_t_ = logN_0_–(log2/T)×t, where N_0_ is the initial NAD concentration (1 mM), N_t_ is the observed NAD concentration at time t, T is the half–life time of exogenous NAD.

## 3. Results and discussions

Degradation of extracellular NAD by *E. coli* cells during whole–cell biocatalysis has been noted recently [5, 7]. This NAD degradation activity should be associated with enzyme outside the cytoplasmic membrane, but no such enzyme has been clearly described in the literature. Because the periplasmic enzyme NadN from *Haemophilus influenzae* had good NAD degradation activity [13], we did sequence alignment and found UshA of *E. coli* shared 29% sequence identity with NadN. Both NadN and UshA have an N–terminal metallophosphatase domain and a C–terminal nucleotidase domain [16]. Interestingly, a previous study found that cell lysates of an *E. coli* BL21 cells transformed with the *ushA* gene expressing plasmid had high hydrolytic activities over a broad spectrum of substrates, such as AMP, ADP, ATP, CMP and CDP, and trace degrading activity for NAD(H) as well as a number of pyrophosphates, but detailed data remained unavailable [9]. We suspected UshA as a candidate periplasmic enzyme responded to NAD degradation.

To demonstrate the function of UshA, we constructed an UshA overexpression strain *E. coli* WL001. The whole–cell NAD degradation activities of WL001, the wild type strain BW25113 and the *ushA* gene deletion strain JW0469 were assayed, and specific activities were 20,000, 199 and 60 U/mg_CDW_, respectively. UshA expression in the three strains was analyzed by SDS–PAGE, a noticeable expression decrease at the molecular weight of 59 kDa was observed in JW0469 compared to BW25113, while UshA was highly overexpressed in WL001 upon induction (Fig. 1). The results confirmed the hypothesis that UshA has a major role in the degradation of extracellular NAD.

**Figure 1.**
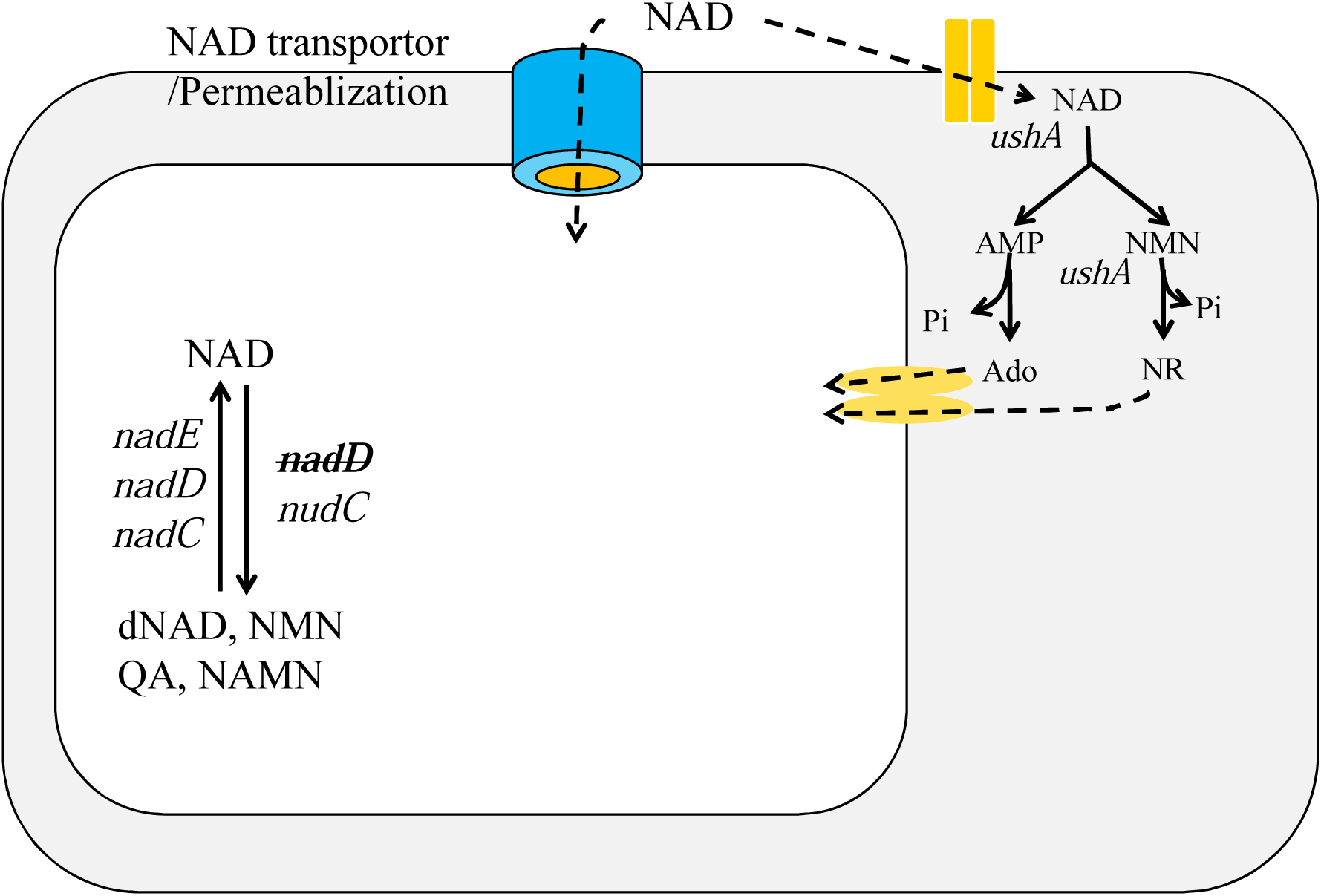
NAD degradation activity of *E. coli* BW25113 and its *ushA::kan* mutant.

To further study the activity of UshA, we purified recombinant UshA from WL001 by His_6_–tag affinity chromatography to near homogeneity (Fig. 1). Analysis of UshA–catalyzed NAD degradation reaction by TLC clearly indicated that NAD was consumed, but there was apparently no accumulation of either AMP or NMN. Instead, the presence of NmR and Ado was evident for those lanes with complete consumption of NAD (Fig. 2A). It was known that NAD can be hydrolyzed to form NMN and AMP by NAD pyrophosphatase [8]. However, TLC results suggested that UshA catalyzed further hydrolysis of NMN and AMP into NmR and Ado, respectively, by 5’–nucleotidase activity (Fig. 2B). Moreover, it seemed that UshA had much higher 5’–nucleotidase activity than pyrophosphatase activity, as both AMP and NMN were not accumulated during the reaction. Therefore, UshA is a promiscuous hydrolytic enzyme which is reminiscent of NadN from *H. influenzae* [13].

**Figure 2.**
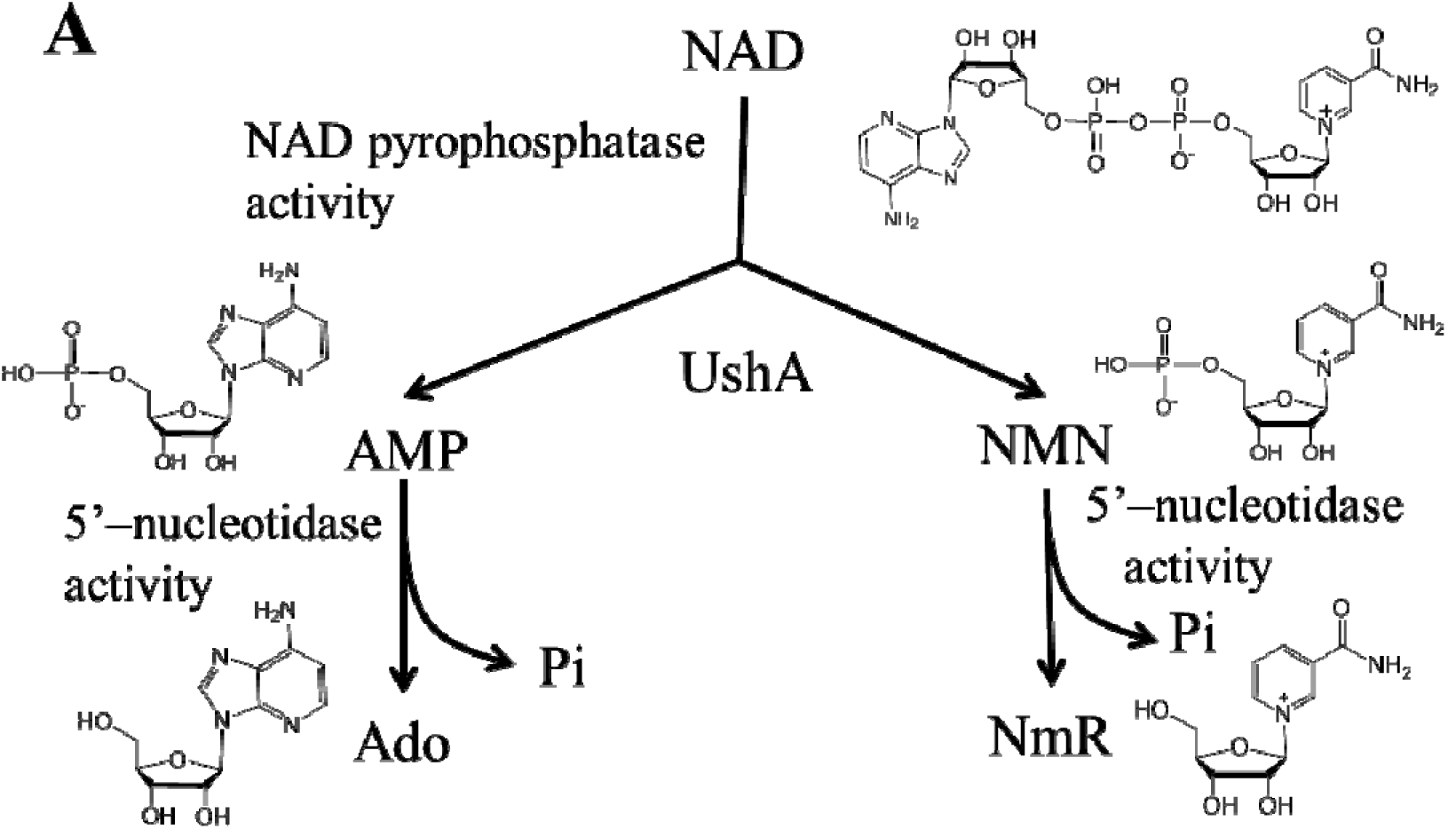
NAD degradation activity of recombinant UshA. Proposed NAD degradation paths by NadN like activity of UshA. Abbreviations: NAD, nicotinamide adenine dinucleotide; AMP, adenosine monophosphate; Ado, adenosine; NMN, nicotinamide mononucleotide; NmR, nicotinamide riboside.

Kinetic parameters of UshA catalysis were determined using NAD, NADH, AMP and NMN, respectively, as substrate (Table 1). While *K*_*m*_ values were close for all of these substrates, *k*_*cat*_ values for AMP and NMN were orders of magnitude higher than those for NAD(H). The *k*_*cat*_*/K*_*m*_ values for AMP and NMN were roughly two orders of magnitude higher than those for NAD(H), indicating that UshA had substantially higher catalytic efficiency for phosphate hydrolysis than pyrophosphate hydrolysis. These data were consistent with results of TLC analysis.

**Table 1.** Kinetic parameters of UshA and other NAD pyrophosphatases.

| Substrate | $K_m$<br>($\mu\text{M}$) | $k_{cat}$<br>($\text{s}^{-1}$) | $k_{cat}/K_m$<br>($\mu\text{M}^{-1} \text{s}^{-1}$) | Enzyme<sup>a</sup> | Reference or<br>resource |
| --- | --- | --- | --- | --- | --- |
| AMP | 1.2±0.1 | 405±4 | 334 | UshA | This study |
| AMP | 1.8 | 372 | 210 | UshA | [9]<sup>b</sup> |
| AMP | 1090 | – | – | NadN | [13] |
| NMN | 11.1±1.3 | 1492±31 | 135 | UshA | This study |
| NMN | 1260 | – | – | NadN | [13] |
| NAD | 5.3±0.6 | 19.3±0.9 | 3.7 | UshA | This study |
| NAD | 430 | – | – | NadN | [13] |
| NAD | 5100±500 | 1.5±0.1 | 2.9×10<sup>-4</sup> | NudC | [8] |
| NAD | 110 | – | – | NudC<sub>Nt</sub> | [18] |
| NADH | 7.0±0.8 | 10.0±0.2 | 1.4 | UshA | This study |
| NADH | 110±20 | 3.8±0.2 | 0.03 | NudC | [8] |

<sup>a</sup> UshA: recombinant *E. coli* UshA; NadN: recombinant *H. influenzae* NadN. NudC: recombinant *E. coli* NudC (synonym of YjaD). NudC<sub>Nt</sub>: wild-type NudC from *Nicotiana tabacum* root.
<sup>b</sup> The reaction condition was the same as this study, with Mg<sup>2+</sup> as the activating cation.

It should point out that the recombinant UshA had the highest NAD pyrophosphatase activity compared to other enzymes collected by BRENDA [17]. The *K*_*m*_ values for NAD (5.3 μM) and NADH (7.0 μM) of UshA were substantially lower, indicating UshA has much stronger binding affinity toward NAD(H) compared to other pyrophosphatases. For NADH degradation, the most efficient pyrophosphatase was *E. coli* NudC with a *k*_*cat*_*/K*_*m*_ value of 0.03 μM^−1^ s^−1^ [8], which was considerably lower than that of UshA (1.4 μM^−1^ s^−1^). Therefore, albeit UshA has higher 5’–nucleotidase activity, it remains the most efficient NAD(H) pyrophosphatase known so far.

We next checked extracellular NAD stability under conditions similar to whole–cell biocatalysis. When NAD was incubated in the presence of permeabilized *E. coli* JW0469 or BW25113 cells, NAD concentration dropped continuously (Fig. 3). Based on these data, the half–life time of NAD in the presence of JW0469 cells was 3.78±0.12 h, which was 3.1–fold of that (1.21±0.08 h) of BW25113 cells. This was consistent with the fact that JW0469 whole cells had only about 30% NAD degradation activity of that of the wild–type strain (*vide ante*). These data clearly demonstrated that the deletion of the *ushA* gene improved extracellular NAD stability, which should be beneficial to cofactor–dependent whole–cell biocatalysts. We noticed that the deletion of the *ushA* gene only reduced NAD degradation activity by 70%, indicating that there were additional gene(s) responsible for periplasmic NAD degradation. Identification of these gene(s) may be interesting for the construction of *E. coli* strains with minimal NAD degradation activity. We recently succeeded in devising *E. coli* NAD–auxotrophic mutants and expanding the dynamic ranges of cellular NAD levels from 0.04– to 9.6–fold of that of wild type cells [6]. If genes responsible for NAD degradation were removed, our capacity for NAD(H) manipulation should reach a new stage, because exogenous NAD(H) could be applied for more robust and precise control on cofactor availability for biocatalysts.

**Figure 3.**
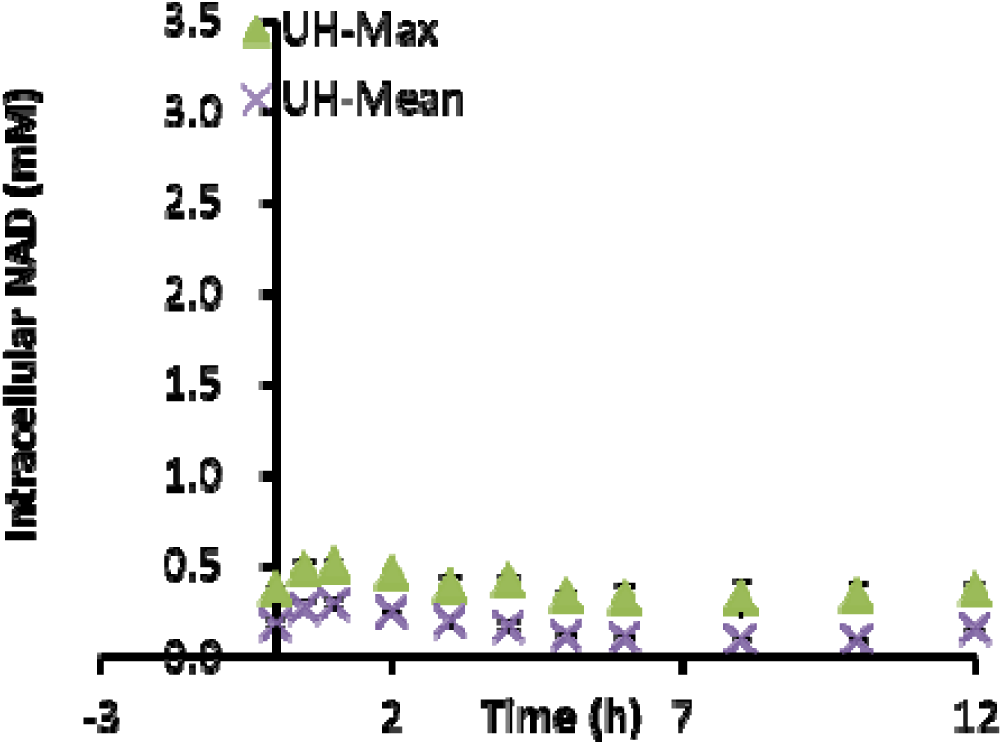
Time course of NAD degradation in the presence of permeabilized *E. coli* cells. Assays were initiated with 1 mM of NAD at 37 °C, pH 7.5. The data represent the averages standard deviations of three independent experiments.

The deletion of the *ushA* gene also led to improved cell growth. Under identical culture conditions in M9 medium JW0469 and BW25113 cells reached a cell density of OD_600_=3.8±0.03 and OD_600_=3.1±0.06, respectively, after 10 h (Fig. 4), indicating the deletion of the *ushA* gene promoted cell growth by 24%.

Our results and previous data [9] indicated that *E. coli* UshA was a promiscuous (pyro)phosphatase enzyme and that its NAD pyrophosphatase activity was noticeably lower than 5’–nucleotidase activity. This activity profile may be advantageous for cells to facilitate utilization of extracellular nutrients under various environments. For biotechnological application such as whole–cell biocatalysis, however, the removal of UshA could improve extracellular NAD stability and cell growth.

## 4. Conclusion

This work demonstrated that UshA was a major periplasmic enzyme for NAD degradation in *E. coli*. The deletion of the *ushA* gene reduced NAD degradation activity by 70% and promoted cell growth. Recombinant UshA showed high NAD(H) pyrophosphatase activity, which significantly enriched our understanding on NAD metabolism. These results should facilitate many applications including designing more robust redox biocatalysts.

## Acknowledgements

This work was supported by the National Basic Research and Development Program of China (No. 2012CB721103) and the State Key Laboratory of Catalysis (R201306).

## References

[1] Holm AK, Blank LM, Oldiges M, Schmid A, Solem C, Jensen PR, et al. Metabolic and transcriptional response to cofactor perturbations in *Escherichia coli*. J Biol Chem. 2010;285:17498–506.

[2] San KY, Bennett GN, Berrios–Rivera SJ, Vadali RV, Yang YT, Horton E, et al. Metabolic Engineering through Cofactor Manipulation and Its Effects on Metabolic Flux Redistribution in Escherichia coli. Metab Eng. 2002;4:182–92.

[3] Berrios–Rivera SJ, Bennett GN, San KY. The effect of increasing NADH availability on the redistribution of metabolic fluxes in Escherichia coli chemostat cultures. Metab Eng. 2002;4:230–7.

[4] Zhang J, Witholt B, Li Z. Coupling of permeabilized microorganisms for efficient enantioselective reduction of ketone with cofactor recycling. Chem Commun (Camb). 2006:398–400.

[5] Zhang W, O’Connor K, Wang DIC, Li Z. Bioreduction with Efficient Recycling of NADPH by Coupled Permeabilized Microorganisms. Appl Environ Microbiol. 2009;75:687–94.

[6] Zhou YJ, Wang L, Yang F, Lin X, Zhang SF, Zhao ZBK. Determining the extremes of the cellular NAD(H) level by using an *Escherichia coli* NAD+–auxotrophic mutant. Appl Environ Microbiol. 2011;77:6133–40.

[7] Zhou YJ, Yang W, Wang L, Zhu Z, Zhang S, Zhao ZK. Engineering NAD+ availability for Escherichia coli whole–cell biocatalysis: a case study for dihydroxyacetone production. Microb Cell Fact. 2013;12:103.

[8] Frick DN, Bessman MJ. Cloning, purification, and properties of a novel NADH pyrophosphatase. Evidence for a nucleotide pyrophosphatase catalytic domain in MutT–like enzymes. J Biol Chem. 1995;270:1529–34.

[9] Alves–Pereira I, Canales J, Cabezas A, Cordero PM, Costas MJ, Cameselle JC. CDP–alcohol hydrolase, a very efficient activity of the 5’–nucleotidase/UDP–sugar hydrolase encoded by the ushA gene of Yersinia intermedia and Escherichia coli. J Bacteriol. 2008;190:6153–61.

[10] Ji D, Wang L, Hou S, Liu W, Wang J, Wang Q, et al. Creation of bioorthogonal redox systems depending on nicotinamide flucytosine dinucleotide. J Am Chem Soc. 2011;133:20857–62.

[11] Baba T, Ara T, Hasegawa M, Takai Y, Okumura Y, Baba M, et al. Construction of Escherichia coli K–12 in–frame, single–gene knockout mutants: the Keio collection. Mol Syst Biol. 2006;2.

[12] van den Ent F, Lowe J. RF cloning: A restriction–free method for inserting target genes into plasmids. J Biochem Bioph Methods. 2006;67:67–74.

[13] Kemmer G, Reilly TJ, Schmidt–Brauns J, Zlotnik GW, Green BA, Fiske MJ, et al. NadN and e (P4) are essential for utilization of NAD and nicotinamide mononucleotide but not nicotinamide riboside in Haemophilus influenzae. J Bacteriol. 2001;183:3974–81.

[14] Lee KS, Song SB, Kim KE, Kim YH, Kim SK, Kho BH, et al. Cloning and characterization of the UDP–sugar hydrolase gene (ushA) of Enterobacter aerogenes IFO 12010. Biochem Biophys Res Commun. 2000;269:526–31.

[15] Baykov AA, Evtushenko OA, Avaeva SM. A malachite green procedure for orthophosphate determination and its use in alkaline phosphatase–based enzyme immunoassay. Anal Biochem. 1988;171:266–70.

[16] Marchler–Bauer A, Lu S, Anderson JB, Chitsaz F, Derbyshire MK, DeWeese–Scott C, et al. CDD: a Conserved Domain Database for the functional annotation of proteins. Nucleic Acids Res. 2011;39:D225–9.

[17] Schomburg I, Chang A, Placzek S, Sohngen C, Rother M, Lang M, et al. BRENDA in 2013: integrated reactions, kinetic data, enzyme function data, improved disease classification: new options and contents in BRENDA. Nucleic Acids Res. 2013;41:D764–72.

[18] Wagner R, Feth F, Wagner KG. The Pyridine–Nucleotide Cycle in Tobacco – Enzyme–Activities for the Recycling of Nad. Planta. 1986;167:226–32.

